# Fish larvae tackle the complex fluid-structure interactions of undulatory swimming with simple actuation

**DOI:** 10.1101/736587

**Authors:** Cees J. Voesenek, Gen Li, Florian T. Muijres, Johan L. van Leeuwen

## Abstract

Most fish swim with body undulations that result from fluid-structure interactions between the fish’s internal tissues and the surrounding water. As just-hatched larvae can swim effectively without a fully-developed brain, we hypothesise that fish larvae tackle the underlying complex physics with simple actuation patterns. To address this hypothesis, we developed a dedicated experimental-numerical approach to calculate the lateral bending moment distributions, which represent the system’s net actuation. The bending moment varies over time and along the fish’s central axis due to muscle actions, passive tissues, inertia, and fluid dynamics. Our 3D analysis of a large dataset of swimming events of larvae from 3 to 12 days after fertilisation shows that these bending moment patterns are not only relatively simple but also strikingly similar throughout early development, and from fast starts to periodic swimming. This suggests also similar muscle activation patterns, allowing fish larvae to produce swimming movements relatively simply, yet effectively, while restructuring their neuromuscular control system.

## Introduction

Swimming is a vital component of the fitness of a fish because fish swim to search for food, hunt prey, escape from predators, migrate and disperse, and manoeuvre through complex environments. Many fish species swim by performing body undulations that result from an interaction between body tissues and the surrounding water [1,2]. Understanding these complex fluid-structure interactions is crucial to gain insight into the mechanics and control of fish swimming [3].

To analyse the fluid-structure interactions during swimming, we need to understand the external fluid mechanics (water), the internal solid mechanics (skin, muscle, skeleton), and their coupling. During swimming, the body of the fish moves through the water, inducing a flow around it [4,5]. The resulting fluid dynamic forces interact with the body tissues via the skin, resulting in a change in deformation. This deformation will change the motion of the surface of the body, which influences the fluid dynamic forces, thus forming a loop of tight coupling between the fluid mechanics and the internal solid mechanics [2,6,7]. This complex two-way fluid-structure interaction creates the typical travelling wave pattern observed in swimming fish [8,9].

The complexity of the physics would suggest that fish need a sophisticated control system to produce swimming motions reliably. Zebrafish (*Danio rerio*, Hamilton 1822) larvae, the subject of this study, would seem to contradict this: they can swim immediately after hatching, at considerable speed and tail beat frequency [3,10,11]. Furthermore, the spinal cord can initiate swimming motions even when severed from the brain [12]. This suggests that a relatively simple system can produce reliable undulatory swimming, despite the non-linear governing physics. Throughout the first days of development, the larvae refine their control of swimming [10,13,14] and improve swimming performance [10,11]. These improvements raise the question: do fish larvae produce swimming differently across early development, and at different swimming speeds and accelerations?

To answer this question, we need insight into the internal mechanics of the axial muscles and passive tissues that actuate the motion. Muscle activation patterns can be measured directly with electromyography, where electrodes are inserted in the muscles to measure potential differences [15–17]. However, this technique may incur considerable changes in swimming behaviour. Especially for fish larvae, it requires them to be paralysed [18] or fixed in place [19], thus changing the fluid-structure interaction that produces the body wave [20]. Furthermore, the resolution along the body is limited by the number of inserted electrodes.

An alternative is an inverse dynamics approach [21], where net forces and moments are calculated from measured kinematics. Hess and Videler [22] used a simplified small-amplitude fluid [23] and internal body model to estimate bending moments along the central axis of saithe from the motion of its centreline. The bending moment is defined for each transversal slice along the body as the sum of the moments produced by the muscles and passive tissues, counteracting the moments due to inertia and water [22,24,25]. Because the muscles are the only component in the system that produce net positive work over a cycle, bending moment distributions are the net actuation of the system, and hint towards properties of the muscle activation patterns.

In this study, we examined bending moment patterns calculated from measured swimming motion across early developmental stages in zebrafish larvae. Previous pioneering studies [22,24] used a small-amplitude model and simplified fluid-dynamics to compute bending moments for only a few cases of periodic swimming in the inertial regime. However, fish larvae use often large undulatory amplitudes and may swim in the intermediate Reynolds regime where inviscid fluid models are invalid. Therefore, we removed these simplifying assumptions and analyse an extensive data set. We used three-dimensional reconstructed kinematics [26], beam theory supporting arbitrarily large deformation amplitudes, and full numerical solutions of the incompressible Navier-Stokes equations to calculate fluid-dynamic forces (Fig 1). With this method, we examined the question: do fish larvae use similar actuation across early development, speeds, and accelerations? We calculated bending moments for 113 swimming sequences of larvae aged between 3 to 12 days-post-fertilisation (dpf). The reconstructed bending moment patterns are qualitatively similar across development, speed, and acceleration. Rather than change the spatiotemporal distribution of bending moments, fish larvae control speed and acceleration with only the amplitude and duration of the bending moment patterns. This suggests that fish larvae retain the same relatively simple actuation to produce swimming across early development, despite complex physics determining the resulting motion.

**Fig 1.**
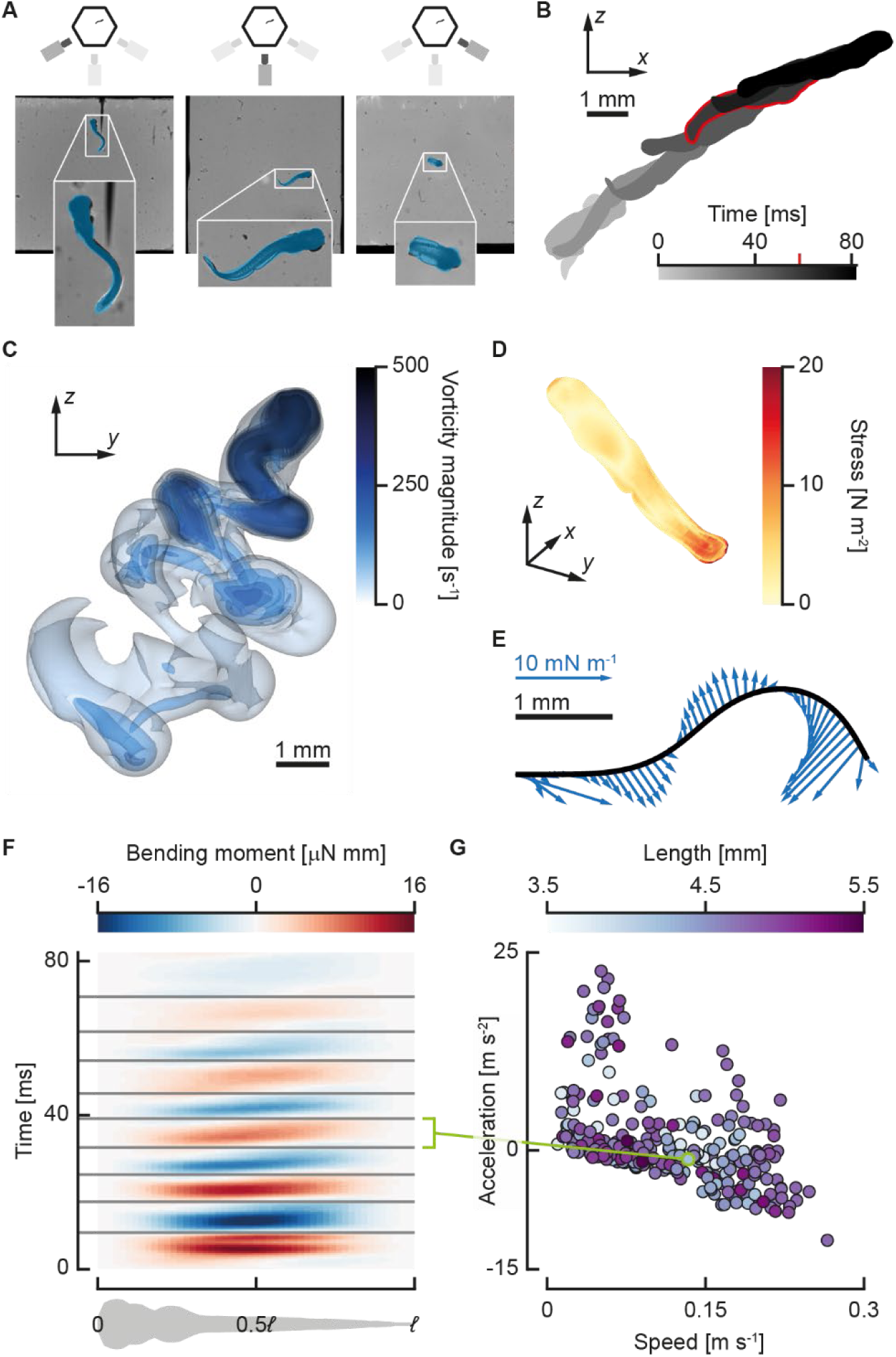
Procedure to calculate bending moments. (A) Larval zebrafish motion is reconstructed from synchronised three-camera high-speed video. Video frames (background) from the three-camera-setup overlaid with projections (blue) of the reconstructed model fish. The legend at the top indicates which camera produced the video frame. (B) Reconstructed three-dimensional motion from the video, projected onto the *x*-*z* plane, the highlighted time instant is shown in (A) and (C–E). (C) Transparent vorticity isosurfaces for the same motion as (B), calculated with computational fluid dynamics (CFD). (D) Total fluid dynamic stress distribution on the skin, the magnitude of the sum of the pressure and shear stress contributions, calculated from the flow field from CFD. (E) Fluid dynamic force distribution transformed to the 2D coordinate system attached to the deformation plane. This distribution is used as input to reconstruct internal moments and forces. (F) Reconstructed bending moment distributions (colour) along the fish (horizontal) and over time (vertical). The horizontal lines separate the half-phases in which the bending moment was divided. The green line links a single half phase to a data point in (G). (G) The mean speed (horizontal), mean acceleration (vertical), and body length (colours) for individual half-beats in the data set (*N* = 285). The green data point corresponds to the highlighted half-beat in (F).

## Results

### Overview of an individual swimming sequence

We performed phase-averaging on a periodic section of a swimming sequence of a 3 dpf zebrafish larva to illustrate how bending moments and bending powers vary along the body during swimming. We selected four half tail-beats (Fig 2A) based on the periodicity of the body curvature. We averaged body curvature, bending moment, kinetic power, and fluid power over these half-beats.

**Fig 2.**
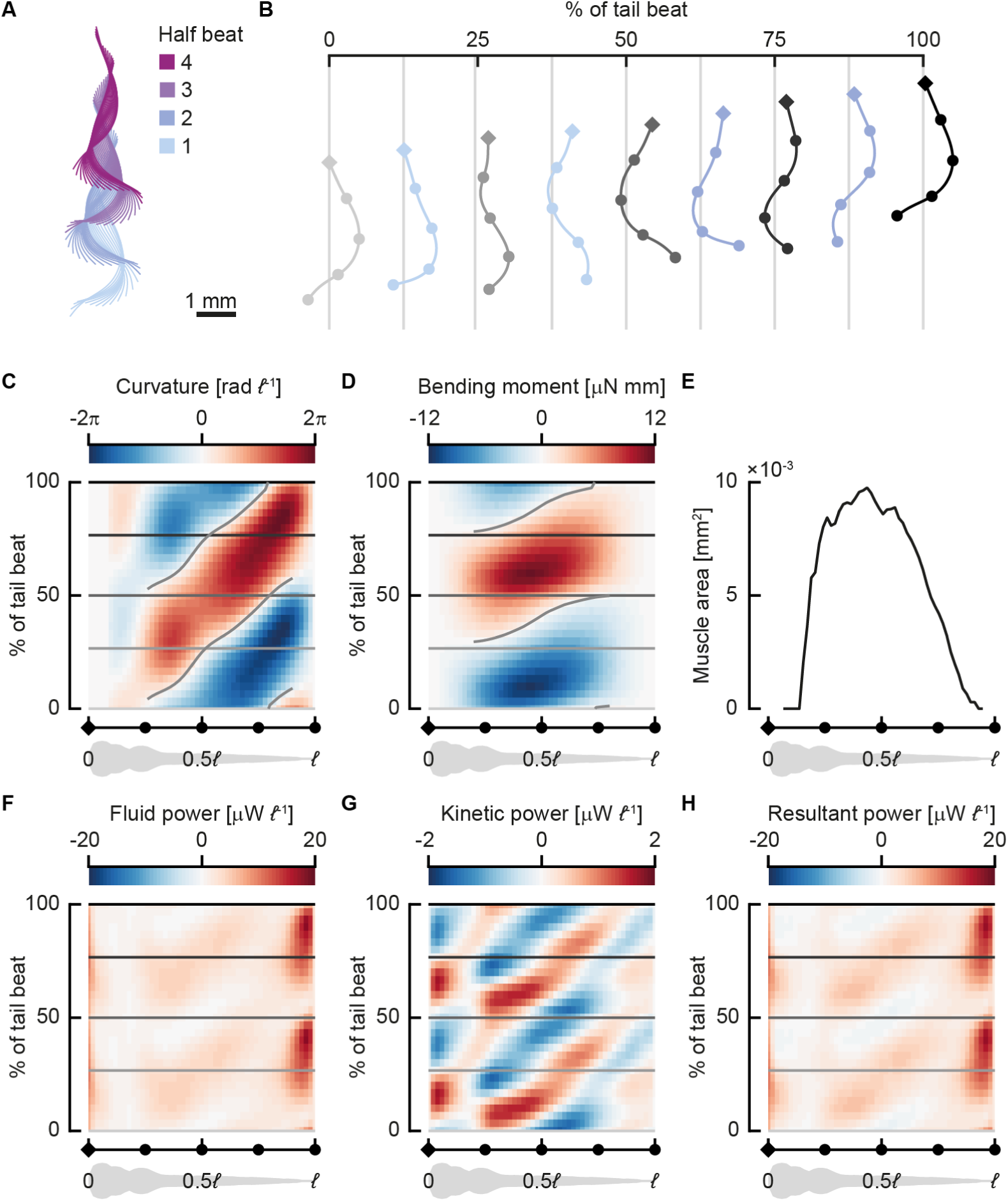
Near-periodic sequence of a 3 days post fertilisation zebrafish larva. The larva swam at 31 *ℓ* s^-1^ with a tail-beat frequency of 69 Hz. (A) Centreline motion throughout the sequence. The colours indicate half-phases. The coordinates were transformed to a best-fit plane through all points along the centreline throughout the motion. (B) Motion during a single full tail-beat (half-beat 1 and 2) of the motion; the grey centrelines correspond to the time points shown with horizontal lines in C,D & F–H. The diamond (head) and dots on the centrelines correspond to points on the *x*-axis for C–H. (C,D,F–H) Heat maps of distributions (colours) along the fish (horizontal) and over the phase over the tail beat (vertical); all quantities are averaged over separate half-beats; “negative” half-beats are mirrored for the curvature and bending moment. (C) Body curvature normalised by body length. (D) Bending moment. (E) Muscle area distribution along the fish. (F) Fluid power per unit body length (power exerted by the fish to move the fluid). (G) Kinetic power per unit body length (rate of change in kinetic energy). (H) Resultant power, the sum of the fluid and kinetic power.

Body curvature (Fig 2B,C) shows a travelling wave pattern behind the stiff head with one positive and one negative peak per cycle. The highest curvatures are reached near the tail, at around 0.8 of the body length (*ℓ*), where body width is relatively small. Curvature waves originate from around 0.25*ℓ*, close to where the most anterior axial muscles are located. They then travel at approximately constant speed (3.3*ℓ* per tail beat) posteriorly, growing in amplitude until close to the tail, and finally dropping to zero amplitude at the tail tip.

Bending moments (Fig 2D) show a positive and a negative peak during swimming, corresponding to the direction of the tail beat, but preceding it in phase along most of the body. The peak amplitude occurs around 0.4*ℓ*, corresponding to the area with the highest muscle cross-section (Fig 2E). Bending moments in the head and tail regions are low due to the free-end boundary conditions, where the bending moment must be zero, the absence of muscle, and in the tail region, the limited cross-sectional area. Like curvature, the bending moment also shows a travelling wave pattern, but its wave speed is more than twice as high as the curvature wave speed (7.1*ℓ* per tail beat).

The power used by the body to move the fluid (Fig 2F) shows a large peak close to the tip of the tail. The motion amplitude is large here (Fig 2A,B), as well as the lateral velocities, therefore fluid forces are large. Since power is the product of velocity and force, most power is expected to be transferred to the fluid here. The kinetic power, defined as the time rate of change in kinetic energy, is smaller in magnitude compared to the fluid power (Fig 2G). The head shows considerable variation in kinetic energy over a tail-beat cycle, owing to its relatively large mass and side-to-side motion. There is a dip in kinetic energy fluctuations in the anterior region of the yolk sac. In the remainder of the body, the kinetic power shows a travelling-wave pattern, caused by the travelling-wave character of the body motion, and hence its speed. The resultant power (Fig 2H), defined as the sum of the fluid and kinetic power, is dominated by the fluid power.

### Swimming effort and vigour

We reconstructed 3D kinematics from 113 video sequences of fast-start responses followed by swimming, calculated flow fields throughout the sequence with CFD, and fitted distributions of internal forces and moments. These swimming sequences hardly contain periodic swimming. To analyse the data despite its aperiodicity, we subdivided it into half-beats based on zero-crossings of the bending moment in the mid-point along the centreline (Fig 1F,G). For each of these 285 half-beats, we calculated the period length, mean speed, mean acceleration of the centre of mass to the next half-beat, and peak (95^th^ percentile) bending moment.

To reduce the number of parameters for the analysis, we identified combinations of parameters with high explanatory capacity. To control swimming, the fish has two main parameters to change the bending moment (see section below): its amplitude and the duration of each half-beat. We define the *swimming effort* as 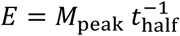 —higher bending moments and shorter periods increase *E*. We fitted a generalised linear model (gamma distribution, log link function) with MATLAB (`fitglm`, R2018b, The Mathworks) and the Statistics and Machine Learning Toolbox (R2018b, The Mathworks). This showed that the swimming effort correlates significantly with the mean resultant power (Fig 3A; *P <* 0.0001), with an exponent of 1.06: close to linear.

**Fig 3.**
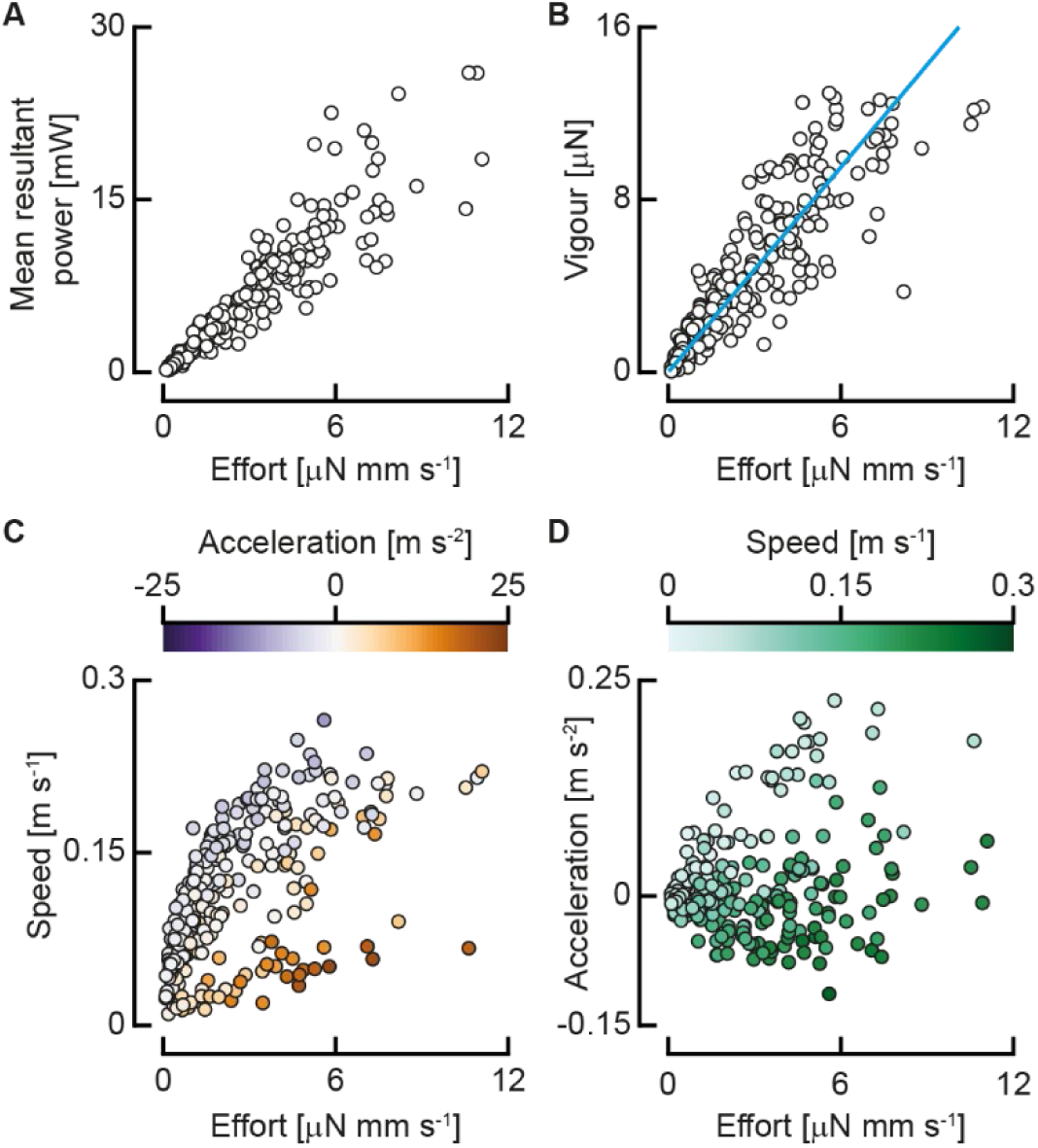
Swimming effort and vigour. (A) The mean resultant power as a function of swimming effort (*N* = 398). (B) Swimming vigour as a function of effort, and the optimal linear fit (*N* = 285). (C) Mean speed as a function of swimming effort, coloured by the mean acceleration (*N* = 285). (D) Mean acceleration as a function of swimming effort, coloured by speed (*N* = 285).

We expect the net propulsive force to scale with the mass of the fish, its acceleration, and its squared speed (from the dynamic pressure). Based on this, we define *swimming vigour* as *V* = *m*(*cv*^2^ + *a*), where *m* is body mass, *v* is swimming speed, and *a* is mean acceleration (i.e. change of speed to the next half-beat per unit of time). The coefficient *c* is calculated with an optimisation algorithm that minimised the sum of squared errors of a linear fit of vigour to effort with total least squares. The fitted value of 517.7 m^-1^ results in a clear trend of vigour as a function of effort (Fig 3B; generalised linear model fit with gamma distribution and log link function: *P* < 0.0001), collapsing the broad clouds of speed and acceleration (Fig 3C,D).

### Bending moment distributions are similar across swimming vigour and development

To assess how bending moment patterns differ across vigour and development (indicated by body length [27]), we compared bending moment patterns normalised by their amplitude. We normalised the bending moment distribution of each half-beat by dividing by the peak bending moment. We then calculated the mean and standard deviation (Fig 4A,B) of the normalised distributions of all half-beats. The standard deviation (Fig 4B) is relatively small, locally peaking at 0.24, caused primarily by variation in the peak phase (Fig 4E,F). For each half-beat, we calculated the mean absolute difference of each point in the distribution to the corresponding point in the mean distribution. The mean of these differences across half-beats is 0.091±0.028—the differences are relatively low, and of similar magnitude across half-beats. Thus, the patterns look similar across different developmental stages and swimming vigour. Note that the peak value is smaller than 1, since the peak location shows variation in both phase and location along the body (Fig 4C–F).

**Fig 4.**
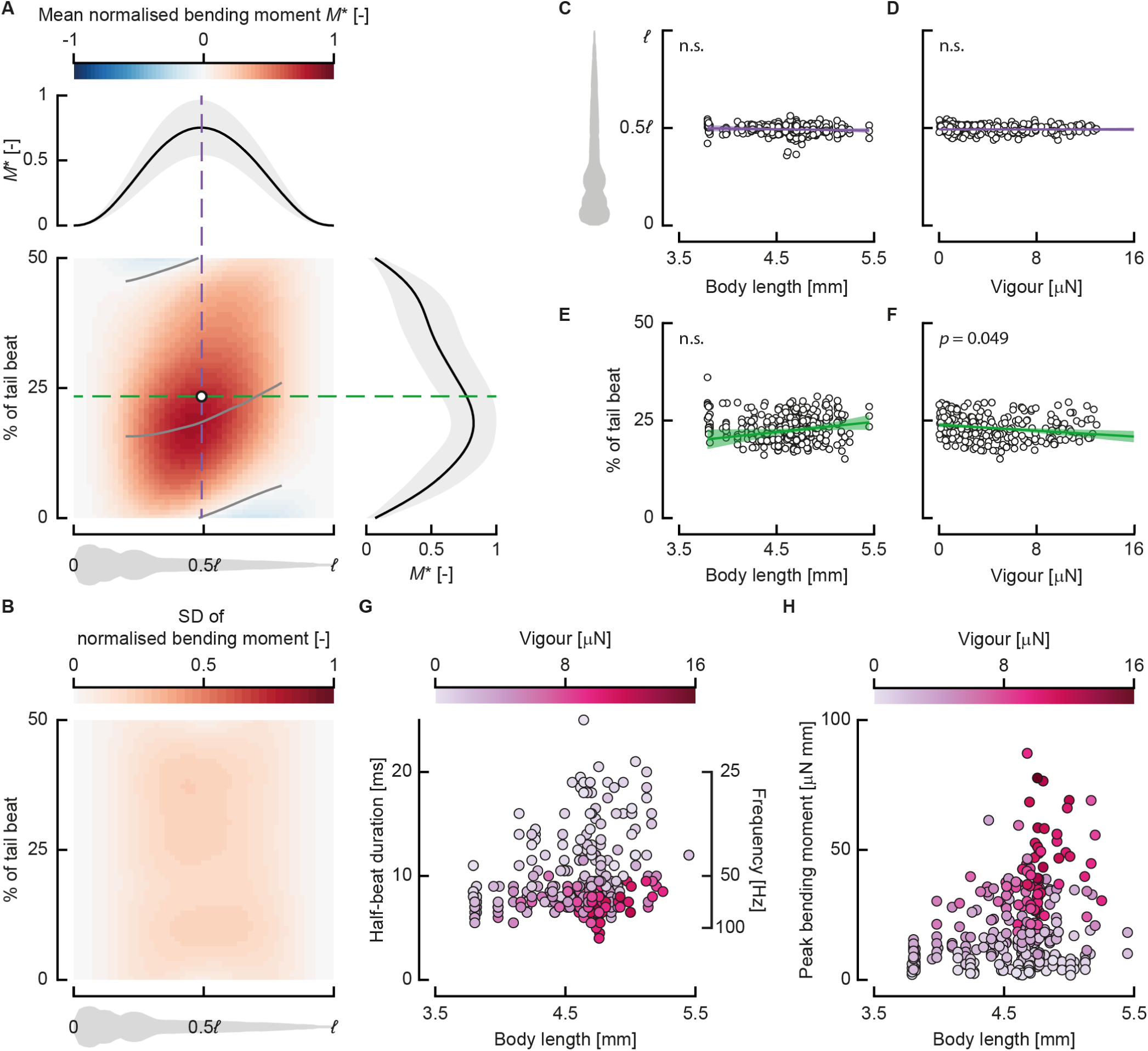
Bending moment patterns are similar across swimming vigour and development. (A) Normalised bending moment pattern along the fish (horizontal) and over normalised time (vertical), averaged over all half-beats (*N* = 398). The dashed lines indicate slices through the pattern in time (green) and location (purple) of the centre of volume of the distribution, shown respectively at the top and right of the heat map, along with their standard deviation. The grey lines over the heat map show the zero and maximum contour line for the middle 60% along the body. (B) The standard deviation (SD) of the normalised bending moment along the fish (horizontal) and over normalised time (vertical) (*N* = 398). (C,*D*) The location along the body of the centre of volume of the bending moment as a function of body length (C, ***N*** = 398) and swimming vigour (D, *N* = 285). (E,F) Normalised time of the centre of volume of the bending moment as a function of body length (E, ***N*** = 398) and swimming vigour (F, ***N*** = 285). (G) Half-beat duration as a function of length (i.e. developmental stage), coloured by swimming vigour (*N* = 285). (H) Peak bending moment as a function of length, coloured by swimming vigour (*N* = 285).

The centre of volume of the individual bending moment patterns (Fig 4C–F) lies around 0.5*ℓ* along the body length and 25% of the tail beat (i.e. 50% of the half-beat). The location along the body varies little across length (i.e. developmental stage) and swimming vigour. The phase (i.e. time relative to the tail-beat duration) shows more variation over length and vigour but shows no clear pattern. We fitted linear models with MATLAB (`fitlm`, R2018b, The Mathworks) with the centre of volume location along the body and in phase as response variable, and the length and vigour as predictors. The slopes for the centre of volume position along the body are not significantly different from 0 for length (*P* = 0.071), vigour (*P* = 0.78) or their interaction (*P* = 0.78). The slopes for the phase of the centre of volume is not significantly different from zero for length (*P* = 0.32) and the interaction between length and vigour (*P* = 0.065), but marginally significant for vigour (*P* = 0.049).

Although the spatiotemporal distributions of the bending moments are similar across lengths (i.e. developmental stage), the duration and amplitude vary (Fig 4G,H). As the fish develop, the range of half-period durations increases (Fig 4G)—young larvae use mostly short durations, while larger larvae use a broad range of durations. The maximum peak bending moment increases substantially over development (Fig 4H). Older fish can generate higher peak bending moments and can reach higher swimming vigour values, but do not always do so.

### Control parameters of swimming vigour

Because the bending moment patterns are similar across swimming styles and developmental stage, the parameters left for controlling swimming vigour are the amplitude of the bending moment and the duration of the tail beat. All experimental points lie on a broad cloud around a curve through the effort landscape, a function of peak bending moment and half-beat duration (Fig 5). In general, high peak bending moments are only produced for tail beats of short duration. As the duration decreases (i.e. frequency increases), the bending moment amplitude decreases. Higher efforts generally lead to higher speeds (Fig 5B), unless the larva is accelerating strongly. Strong accelerations are mostly found with slow-swimming larvae using short half-beat durations and high peak bending moments (Fig 5A). For high-effort tail beats, the larvae are generally either swimming fast, or accelerating: high-effort, low-speed tail beats show high accelerations, while high-effort, low acceleration tail beats show high speeds. Swimming vigour tends to increase with increasing effort (Fig 5C, see also Fig 3B).

**Fig 5.**
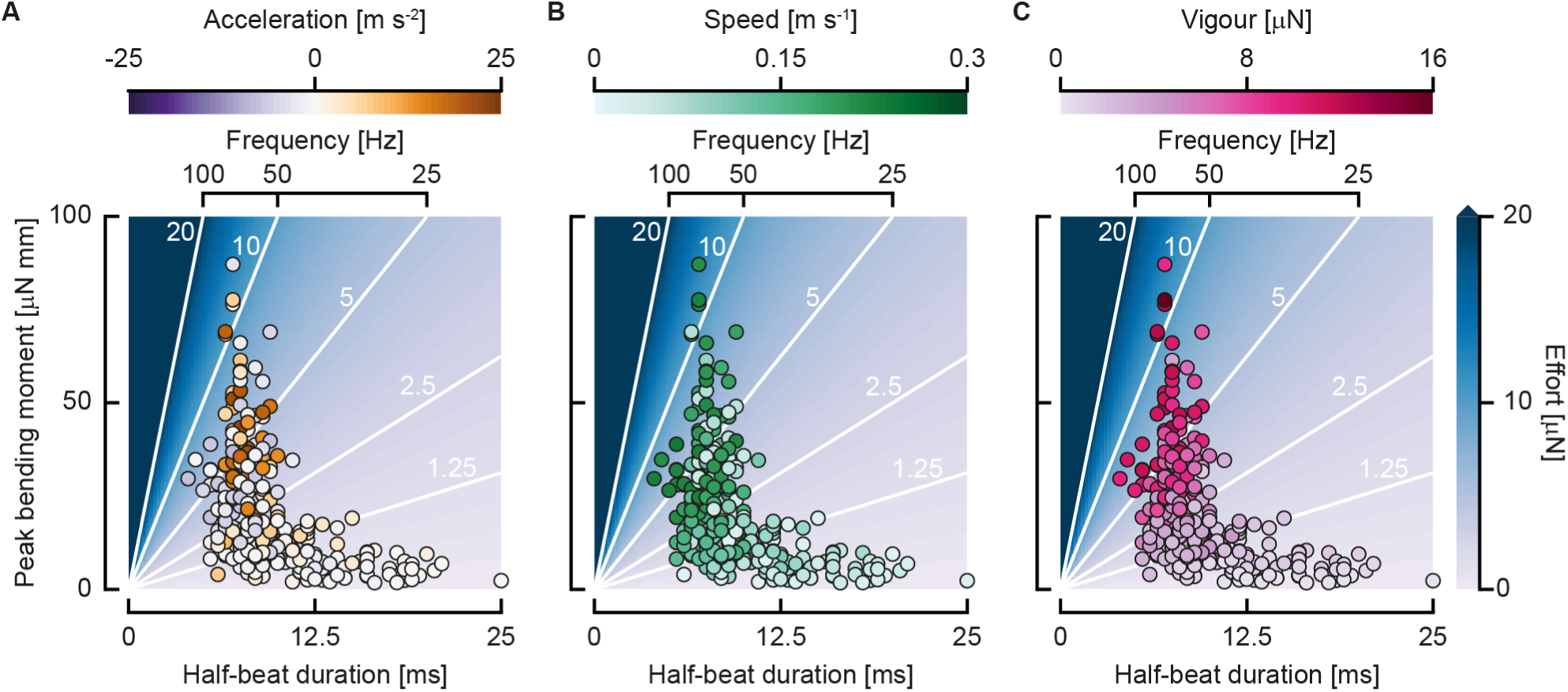
Swimming control parameters. (A,B,C) Individual half-beats in the duration – peak bending moment landscape, the coloured background with white contour lines shows the swimming effort (*N* = 285). (A) Dots coloured by acceleration. (B) Dots coloured by speed. (C) Dots coloured by swimming vigour.

## Discussion

In this study, we analysed the actuation used during swimming of developing zebrafish larvae by calculating bending moment patterns with inverse dynamics. We found that larvae use similar bending moment patterns across development. They adjust their swimming vigour, a combination of speed and acceleration, by changing the peak bending moment and tail-beat duration. At higher speeds and accelerations, the larvae produce the required fluid-dynamic forces by increasing bending moment amplitude and/or decreasing tail-beat duration.

Previous inverse-dynamics approaches for the internal mechanics of swimming used simplified models for both the fluid and the structure. The structure was modelled with linear bending theory, assuming small deformations of the centreline [22,24]. The effects of a large-amplitude correction to these was expected to be small for adult fish that swim periodically [28]. However, zebrafish larvae beat their tails often at >90° with the head [11], violating the small-amplitude assumption. The beam theory underlying our bending moment calculations allows arbitrarily large deformation, under the assumption of pure bending. However, our beam model ignores the effect of shear deformation that is expected to occur close to the medial plane [29], but we expect it to be of small influence on the bending moments due to its proximity to the axis and hence small moment arm.

In addition to the large-amplitude beam model, we used 3D CFD to calculate fluid-dynamic forces, dropping previous assumptions of inviscid flow [22–24,30] and the necessity to model the boundary layer separately [28]. The assumption of inviscid flow does not hold for fish swimming in the intermediate regime [3,31]—full solution of the Navier-Stokes equations is necessary to obtain sufficiently accurate fluid-dynamic force distributions. The intermediate Reynolds number of the zebrafish larvae allows us to solve the Navier-Stokes equations accurately, without requiring turbulence modelling [31].

In our analysis of swimming sequences across development, we do not assume periodicity. Periodic motion is a common assumption in the analysis of fish swimming [11,32]. For zebrafish, cyclic swimming occurs most often after a fast start, and rarely spontaneously [10,33], and generally only for a few tail beats. To analyse aperiodic motion, we subdivided each swimming sequence in tail beats based on zero-crossings of the bending moment in the middle of the body. During aperiodic swimming, the mean speed varies between successive tail beats. For this reason, we define a parameter “swimming vigour”, that combines the effects of acceleration and swimming speed as *V* = *m(cv*^2^ + *a)*. This approach for analysing aperiodic swimming could be of general use in swimming research. The subdivision in half-beats can also be done with quantities other than the bending moment, for example body curvature. This enables similar analyses of aperiodic swimming from pure kinematics without inverse dynamics.

We define the swimming effort exerted by the fish based on the amplitude of the bending moment and the duration of the tail-beat, as 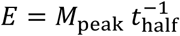. This quantity correlates with resultant power, indicating that it is indeed an indicator of amount of effort exerted by the larva (Fig 3A). The speed and acceleration fall on broad clouds as a function of effort (Fig 3C,D), since the required power depends on their combination, rather than their individual values. The effort-speed landscape (Fig 3C) shows a two-pronged distribution, one branch showing high effort but low speed, and the other, broader branch showing increased effort with speed. This distribution is mainly explained by the acceleration, showing high values in the lower branch—fish only accelerate strongly from low speeds and use high effort to do so. This is reflected in the effort-acceleration landscape (Fig 3D), low (including negative) acceleration are found at high speeds, and *vice versa*.

When speed and acceleration are combined into the swimming vigour, these clouds collapse more closely to a trend line (Fig 3B). Variation in this trend may be partly caused by turning behaviour and contributions of the pectoral fins. We can estimate the relative contribution of the speed and acceleration to the swimming vigour, giving an indication of their relative cost. If we assume force production to maintain speed and to accelerate are equally costly, we can estimate a drag coefficient from the coefficient in the vigour equation as 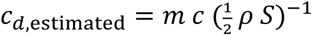, where *ρ* is the fluid density, and *S* is the wetted area. Its value is 0.061, which is considerably lower than the value of 0.26 calculated from a previous CFD study on larval zebrafish [34]. This means our equal-cost assumption does not hold: the contribution of the speed term is relatively low compared to the acceleration term. Since swimming vigour correlates with swimming effort, this indicates that acceleration is more costly to achieve compared to maintaining swimming speed—the larvae need to invest more effort to produce force to accelerate than to swim steadily.

Most of the resultant power produced by the fish is used to increase the energy in the fluid, rather than the kinetic energy of the body (Fig 2H). The energy spent on the water is likely lost on lateral velocity: larvae swim at high Strouhal number, associated with large tail-beat amplitudes and relatively high energy consumption [11,35]. Most of this fluid power is produced at the tail, where the largest fluid-dynamic forces are produced [34], even though no muscles are present here. This suggests a transfer mechanism by passive tissues from the muscles to the tail [36–38].

The bending moment does not correspond directly to muscle action, as it also includes the effects of passive structures inside the body [22]. We do not know the contribution of the muscles to the total bending moment, nor the specific distribution of stresses inside the body. Cheng *et al.* [25] modelled the elastic and visco-elastic properties of the passive tissue, and thus estimated the contribution of the muscle bending moment. The amplitude of the muscle bending moment was found to be higher than the overall bending moment, while the wave speed was found to be lower. However, the overall dynamics look reasonably similar. If we assume similar distributions of passive tissues inside the fish across the considered developmental stages [27], similar total bending moment patterns will require a similar muscle contribution. Furthermore, the difference in amplitude between similar bending moment patterns must originate from the muscle moment, since it is the only net source of power in the system – the work done by the fluid and passive tissues indirectly comes from the muscles.

We found that the bending moments follow a similar pattern across development and swimming vigour (i.e. speed and acceleration). The only significant coefficient in the linear models is the phase of the centre of volume of the bending moment patterns as a function of swimming vigour (Fig 4F), but the effect is limited. More vigorously swimming fish generate the peak bending moment slightly earlier in the half-beat. The mean pattern looks qualitatively similar to earlier calculations done for adult fish [24]. It is a single-peaked distribution, with the maximum around the bulk of the muscle (Fig 2E, Fig 4A), and a fast-travelling wave character. Muscle electromyograms (EMG) done on paralysed zebrafish also looked similar to adult activation patterns [18]. This suggests that this simple pattern of bending moments is common to fish across species and developmental stage. Even though fish larvae swim in the intermediate regime [3], and adult fish often swim in the inertial regime [39], the differences in fluid dynamics do not seem to require fundamentally different bending moment patterns.

Since the bending moments look similar along the body and over the phase for each half-beat, the two parameters left for the larvae to adjust for each half-beat are its duration and the amplitude of the bending moment. Young larvae use a relatively narrow range of amplitudes (Fig 4H) and durations (Fig 4G), which broadens as the fish develop. Older larvae are able to generate higher peak bending moments, likely correlated to development of their muscle system [40]. Furthermore, older larvae use a broader range of tail beat durations than young larvae, suggesting that older larvae have more freedom to control their swimming vigour.

Swimming kinematics emerge from simple bending moment patterns. These patterns presumably stem from simple muscle activation input—their quantification is an interesting avenue for future research. The arrangement and properties of the muscles, passive tissues and propulsive surface causes simple inputs to translate into complex kinematics and flow fields. This has profound consequences for larvae that need to swim to survive [41]. Straight from the egg, they can produce swimming behaviour to escape threats, despite relatively limited neural processing capacity. This concept of designing passive systems to allow complex systems to be actuated simply is of broad interest in engineering and biology [42–44]. Because bony fish larvae are similar in morphology across many species [45], we expect that these results are relevant for bony fish in general. The simple actuation solution may have been instrumental in the evolution and adaptive radiation of bony fish (with more than 30,000 extant species).

## Methods

An in-depth mathematical treatment of the methods is given in the Supplemental Information.

### Reconstructing 3D motion from multi-camera high-speed video

We made high-speed video recordings of fast starts of three separate batches of 50 zebrafish larvae from 3–12 days post fertilisation (dpf). The camera setup was identical to the setup described in Voesenek *et al.* [26], with three synchronised high-speed video cameras, recording free-swimming larvae at 2000 frames per second. To reconstruct the swimming kinematics from the recorded high-speed video, we used in-house developed automated 3D tracking software [26] in MATLAB (R2013a, The Mathworks). For every time point in a multi-camera video sequence, the software calculates the best fit for the larva’s 3D position, orientation and body curvature to the video frames. These parameters are then used to calculate the position of the larva’s central axis and the motion of its outer surface (Fig 1A,B).

### Subdividing motion

We calculated phase-averaged quantities for an individual swimming sequence to look in detail at the generated bending moments and powers. We determined whether a (subset of a) sequence is periodic with a similar approach to Van Leeuwen *et al.* [11]. For every possible subset of a swimming sequence, we calculated the sum of absolute difference with a time-shifted version of the curvature, similar to an autocorrelation. We then calculated extrema in this function – if extrema are detected, their maximum value determines the “periodicity” of the sequence. We then selected the longest possible subsequence that has a periodicity value higher than a threshold of 35 – this is a swimming sequence for a 3 dpf fish. We divided this sequence in half-phases based on peaks in the body angle [11,26]. The curvature, bending moment, fluid power, kinetic power, and resultant power were then phase-averaged based on these subdivisions.

Most of the swimming of larval zebrafish is aperiodic, but there is an alternating pattern in the bending moments. For this reason, we analysed swimming per half-beat, based on the bending moment. We found the zero crossings of the bending moment at 0.5*ℓ*. Since some of these points are related to noise, we evaluated every possible permutation of zero crossings per sequence on several criteria with a custom MATLAB (R2018b, The Mathworks) program. We eliminated zero-crossings with neighbouring sections with an amplitude of less than 5% of the peak half-beat amplitude in the sequence, as they are most probably noise. We required more than three zero crossings to have at least two half-beats to be able to calculate a mean acceleration. Extreme values in each half-beat should alternate direction to eliminate noisy zero crossings: the larva beats its tail left and right, so therefore bending moment must alternate. Finally, we eliminated half-beats with a duration shorter than 2.5 ms (equivalent to 200 Hz tail-beat frequency) – the maximum tail-beat frequency observed for zebrafish larvae is 95 Hz [11]. From all permutations that met the criteria, we selected the permutation with the smallest standard deviation in half-period length across the sequence. This left the longest possible, least noisy sequence of half-beats for every swimming bout.

Out of 113 swimming sequences, we selected 398 half-beats with this procedure. For each of these half-beats, we calculated the duration, mean speed, and peak bending moment. We determined the mean acceleration by calculating the difference in mean speed between the following and current half-beat. Since we could not calculate mean acceleration for the last half-beat in each sequence, 285 half-beats remained for which we computed all quantities.

### Calculating fluid-force distributions

To calculate fluid-force distributions, we used the immersed boundary method Navier-Stokes solver IBAMR [46]. We converted the tracked video data into a three-dimensional point cloud model in the fluid solver. We exclusively used swimming sequences where the larvae start from rest in quiescent water, so we do not need to consider history in the wake. The solver time step was much smaller than the time step between video frames, so we interpolated the reconstructed kinematics with a quintic spline [47]. Using this interpolated state, we updated the location of the point cloud representing the surface of the fish. This resulted in a three-dimensional velocity and pressure field at every point in time (Fig 1C). To verify the accuracy of the method, we compared reconstructed bending moments from IBAMR to an experimentally validated CFD solver [31,34,48], showing only small differences, see the Supplemental Information.

We extracted force distributions by interpolating the pressure and velocity gradient tensor components to the centre of each face of an offset triangulated representation of the fish surface. We then integrated these values into contributions to the pressure force and the shear force at every face of the surface (Fig 1D). By further integration, we calculated the force at every point along the centreline in a coordinate system attached to the larva’s head (Fig 1E).

### Calculating bending moments

To calculate bending moments, we represented the fish by its central axis only. Effects of muscles, spine, and other tissues were combined for every transversal slice along this axis. This simplification allowed us to describe the fish as a non-linear, one-dimensional beam in two-dimensional space. We derived the equations of motion for this beam (see Supplemental Information) in an accelerating and rotating coordinate system attached to the fish’s head [49].

We obtained the motion of the fish from the tracked video, and the fluid forces from the fluid model. This left the normal forces, shear forces, and the bending moment as the only unknowns in the equations. We described the distributions of these unknowns with a quintic spline [47] with uniformly spaced control points along the axis.

To determine the control point values of the normal force, shear force and bending moment, we minimised the residuals of the equations of motion. For every trial combination of control points, we calculated the residuals of equations at all points along the fish. The squared sum of these normalised residuals was minimised with a Levenberg-Marquardt algorithm [50] to obtain the best-fitting control point values that meet the boundary conditions for both free ends (internal forces and moments are zero). When the residuals of the equations are equal to zero, the optimised distributions satisfy the governing equations and boundary conditions exactly. Therefore, this procedure ensured that the computed internal force and moment distributions (Fig 1F) were as close to physically valid as possible within the measurement error. From the motion of the centreline and fluid dynamic forces, we derived a local resultant power.

This optimisation procedure was validated with reference data obtained by integrating the equations of motion with a known external force distribution and internal moment distribution. We then reconstructed the bending moments, shear force, and normal force from the integrated motion and the prescribed external force distribution, resulting in near-identical values (see Supplemental Information).

These data were also used to calculate the resultant power from the fluid-dynamic forces and the changes in kinetic energy. Note that we cannot calculate a meaningful value for the internal power, because we do not separate passive and active effects. The net power when combining these effects does not correspond to the actual power consumption of the fish, as the passive and active components might partly compensate each other.

### Calculating muscle cross-sectional area

We performed micro-computed-tomography (μCT) images of a 3 dpf zebrafish larva at the TOMCAT beamline at the Paul Scherrer Institut [51]. The larva was fixed in Bouin’s solution and stained with phosphotungstic acid (PTA). The complete fish was imaged by stitching three scans with a resolution of 650×650×650 nm per voxel. From these data, a centreline was extracted by finding the centre of area of each slice, segmented with simple thresholding. Finally, the muscle area was manually digitised in 51 planes, for which the image data was interpolated in a plane perpendicular to the centreline.

## Acknowledgements

We thank the staff of the Carus animal facilities for providing the zebrafish larvae; Remco Pieters for building the camera setup; Stefan de Vilder from SDVISION for providing the pco.dimax HS4 camera; Herman ten Berge from Acal BFi Nederland for providing the Photron FASTCAM SA5 camera; Karen Léon-Kloosterziel, Remco Pieters, and Henk Schipper for their assistance during the experiments; Wouter van Veen for helpful discussion on CFD; the Paul Scherrer Institut for use of the TOMCAT beamline at the Swiss Light Source, and Rajmund Mokso for providing guidance with the CT-scans. This work was supported by grants from the Netherlands Organisation for Scientific Research (Nederlandse Organisatie voor Wetenschappelijk Onderzoek; NWO/ALW-824-15-001 to J.L.v.L. and NWO/VENI-863-14-007 to F.T.M) and the Japan Society for the Promotion of Science (JP17K17641 to G.L.).

## Author contributions

Conceptualization: CJV, JLvL

Data curation: CJV

Formal analysis: CJV, JLvL

Funding acquisition: JLvL

Investigation: CJV, JLvL, GL, FTM

Methodology: CJV, JLvL, GL, FTM

Project administration: JLvL, CJV

Resources: JLvL

Software: CJV, GL, JLvL

Supervision: JLvL, FTM

Validation: CJV, GL, JLvL

Visualization: CJV

Writing– original draft: CJV, JLvL

Writing– review & editing: CJV, JLvL, FTM, GL

## Supplementary Information

In this Supplemental Information, we provide the detailed mathematical background of the methods used in the article “Fish larvae tackle the complex fluid-structure interactions of undulatory swimming with simple actuation”. In addition, we show the results of the validation performed on these methods.

### 1 Equations of motion

In this study, we calculated internal forces and moments for swimming zebrafish larvae. The three-dimensional motion of the larvae was obtained from multi-camera high-speed video with an automated tracking method^1^. From this motion, we calculated the internal forces and moments by modelling the fish as a bending ‘beam’. In this section, we show the derivation of the equations of motion for this large-deformation beam representation of the fish.

#### 1.1 Deriving the equations of motion

We model the fish as a beam with varying cross-sections, undergoing arbitrarily large deformation. Plane cross-sections are assumed to remain plane and perpendicular to the neutral line (no shear deformation), but axial deformation is allowed. Although the motion we tracked from the video is three-dimensional, we assume that the fish deforms in a single plane. Therefore, we can use a two-dimensional beam model to represent the deformation of the fish, under a suitable coordinate transformation. In summary, we model the fish as a beam undergoing large bending deformations in two dimensions.

We describe the deformation of the beam with the displacement of each infinitesimal beam element with respect to the reference configuration (Fig. 1A). It is defined as a function ξ(*s, t*) = *ξ*(*s, t*), *η*(*s, t*) of a parameter *s* along the length of the beam, and the time *t*. The position of the central axis at each point *s* is given by:

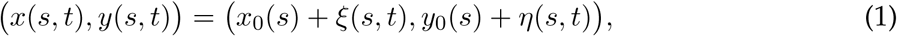

where (*x*_0_(*s*), *y*_0_(*s*)) is the reference configuration of the beam. We define the reference configuration as a straight beam aligned with the positive *x*-axis, so the position becomes:

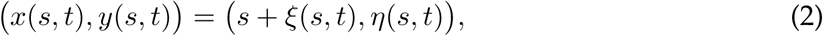

This displacement results in a local deformation angle *θ*(*s, t*) for each beam element (Fig. 1A). It can be calculated from the displacements with:

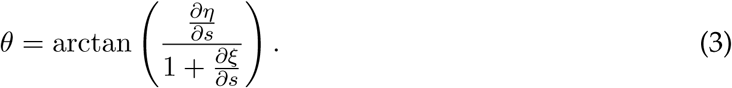

**Fig. 1.**
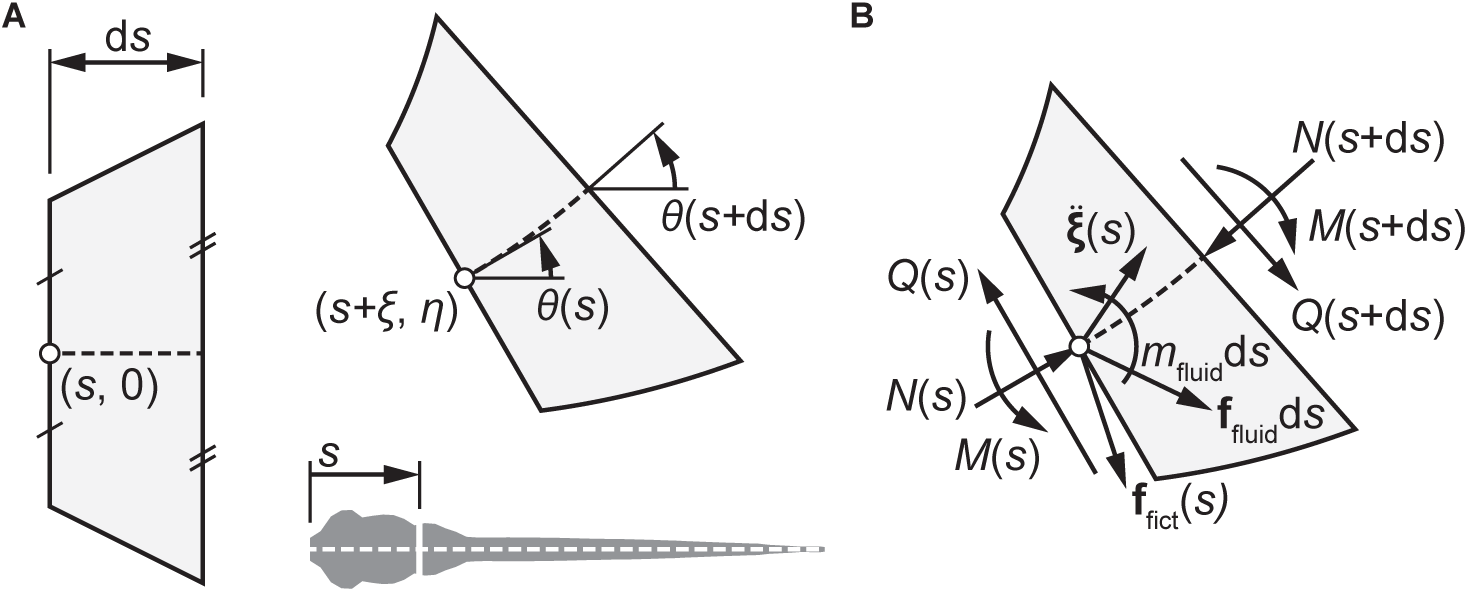
Beam representation of the fish body for computation of bending moments. (A) Geometric definitions of a beam element d*s* at a position *s* along the body, in reference (left) and deformed (right) configuration. We consider an infinitesimal element at (*s*, 0) in the reference configuration, that is displaced by (*ξ, η*), and rotated by an angle *θ*. (B) Free body diagram of a beam element d*s*, with axial force *N*, shear force *Q*, bending moment *M*, net fictitious force distribution **f** _fict_, net fluid force distribution **f** _fluid_, net in-plane fluid moment distribution *m*_fluid_, and acceleration 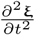.

To derive the equations of motion, we consider a free body diagram of an infinitesimal beam element of length d*s* (Fig. 1B). The force balance in *x*- and *y*-direction for a beam element d*s* can be expressed as:

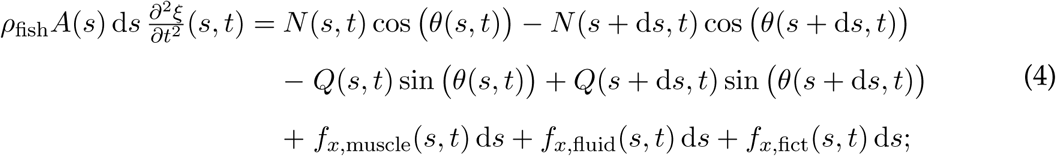

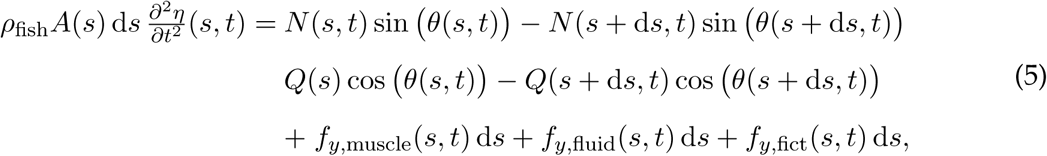

where *ρ*_fish_ is the beam density, *A* is the beam cross-sectional area, *N* is the internal normal force, *Q* is the internal shear force, *f*_*x*,muscle_, *f*_*y*,muscle_ are the muscle forces in *x*- and *y*-direction, *f*_*x*,fluid_, *f*_*y*,fluid_ are the external fluid forces in *x*- and *y*-direction, and *f*_*x*,fict_, *f*_*y*,fict_ are the fictitious forces in *x*- and *y*-direction (from the non-inertial reference frame, see section 1.2).

The moment balance (counter-clockwise positive) about the point on the neutral line at *s* + d*s* is given by:

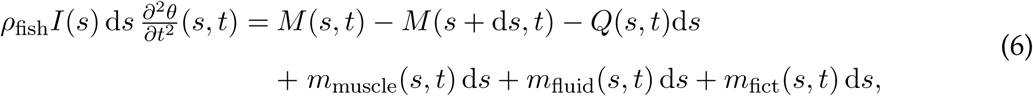

where *I* is the distribution of the second moment of area, *M* is the internal moment, *m*_muscle_ is the muscle moment, *m*_fluid_ is the external fluid moment, and *m*_fict_ is the fictitious moment (see section 1.2).

Dividing equations 4, 5, and 6 by the infinitesimal length d*s*, applying the definition of a derivative 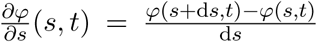, and dropping the explicit *f* (*s, t*) notation, yields the equations of motion:

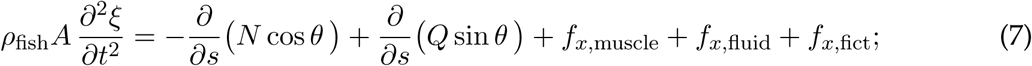

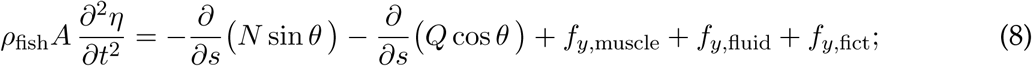

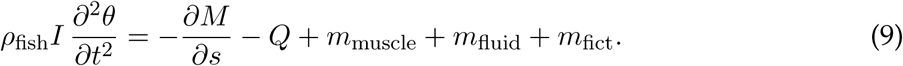

#### 1.2 Fictitious forces

We reconstructed the motion of the fish from video in three-dimensional space, but described the equations of motion in a two-dimensional plane. However, in the video-tracking method, we assumed that the fish deforms in a single plane. Hence, we can create a coordinate system aligned to this plane and obtain the equations in two dimensions only. We define this head reference frame as fixed to the snout of the fish and rotating along with the stiff head region in the deformation plane. Any point 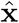 in world coordinates (denoted with a circumflex) can be expressed in the fish coordinate system at time *t* as:

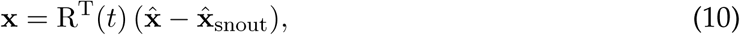

where R(*t*) is the time-dependent rotation matrix expressing the orientation of the snout and 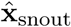 is the position of the snout.

When we transform the motion to the non-inertial fish reference frame, additional equation terms accounting for the effect of the translation and rotation of the frame must be considered. These additional acceleration terms for any point **x** in the rotating reference frame are given by^2^:

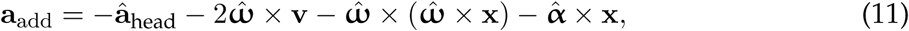

where â_head_ is the acceleration of the origin, 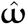 is the angular velocity of the rotating coordinate system, **v** is the velocity of the considered point, and 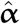 is the angular acceleration of the rotating coordinate system. Note that the quantities 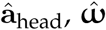, and 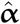 are calculated with respect to the world reference frame, but expressed in the basis vectors of the moving reference frame. The position **x** and velocity **v** are expressed with respect to the moving reference frame.

These accelerations can be considered as an additional ‘fictitious’ external force distribution in the moving reference frame. This force distributions is given by

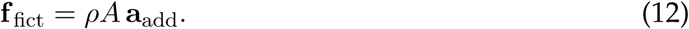

These fictitious forces are added to the fluid-dynamic forces to calculate the total external force distribution acting on each beam element.

### 2 Calculating fluid forces from kinematics

The equations of motion include an external force distribution, produced by the water on the skin. Since this is exceedingly difficult to measure directly and non-invasively, we modelled the fluid dynamics numerically. We used two independent computational fluid dynamics (CFD) methods to calculate fluid-dynamic forces. We used an experimentally validated method to validate the second method to assess its accuracy when calculating internal forces and moments.

We performed computational fluid dynamics using a Navier-Stokes solver based on over-set meshes^3,4,5^, coupled to a body dynamics solver to simulate free swimming. Simulations were performed with swimming kinematics based on a travelling wave with a known curvature amplitude at a frequency of 50 Hz. The same motion was used in a second, independent Navier-Stokes solver based on the immersed boundary method, the open-source code IBAMR^6^.

The Navier-Stokes equations were solved on a rectangular domain, with extents determined by the bounding box around the complete motion with an additional margin of 2 fish lengths. The immersed boundary solver used an adaptive mesh refinement approach, in which the computational mesh can be locally refined depending on the flow conditions. In our case, the mesh consisted of four levels of refinement. Each level was a simple rectangular Cartesian mesh with 4 times the number of subdivision in all dimensions compared to the coarser level. The choice of mesh refinement level depended on the local value of the vorticity, we chose thresholds of 1, 25, and 250 s^−1^ to switch to the second, third, and fourth refinement level respectively. We used a fixed time step of 0.5 µs (see section 6.2), where the CFL-number is always much smaller than 1. We saved the fluid solution every 0.25 ms—at these points we reconstruct the internal forces and moments.

The surface of the fish was described as a cloud of Lagrangian points, moving over the Eulerian fluid solution mesh. The motion of these points was prescribed based on quintic spline interpolation^7^ of the tracked kinematics, with a custom-developed add-on to IBAMR. The density of the point cloud was chosen such that the mean distance between the points is 0.75× the smallest mesh level. This ensured that each cell inside the fish body will have at least one point in it, and not much more.

The resulting flow fields were post-processed to extract the fluid force distribution on the skin of the fish with a custom Python 3 program. In this program, we interpolated^8^ the pressure and velocity gradients to a triangulated surface slightly offset from the fish skin. These were then used to calculate the local stress on each triangular face as:

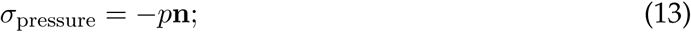

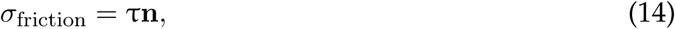

where *p* is the pressure, τ is the shear stress tensor, and **n** is the outward facing normal of the face. Under the assumption of a Newtonian fluid, the shear stress tensor is defined as:

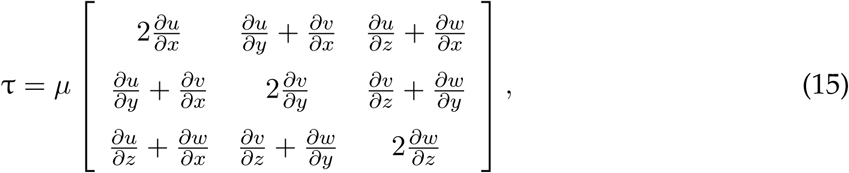

where *µ* is the dynamic viscosity, and *u, v, w* are the velocity components in respectively *x*-, *y*-, and *z*-direction. These surface stress distributions were then grouped into segments along the fish to calculate the local net fluid force in the moving reference frame.

### 3 Reconstructing internal forces and moments with inverse dynamics

We reconstructed bending moments from the motion of the fish and its simulated external fluid force distribution, an approach commonly called inverse dynamics. This section describes the optimisation procedure we used to reconstruct the internal forces and moments.

In our inverse dynamics approach, we cannot separate the effects of the active and passive tissues inside the fish: the internal forces and moments that we compute include the effects of both. Considering this, the moment equation becomes:

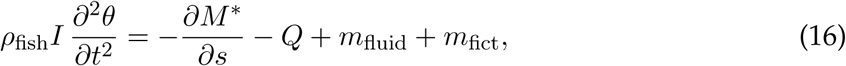

where 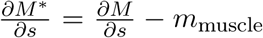. This combined moment is what we reconstruct from the motion with our optimisation procedure. From here onwards, we drop the asterisk notation and refer to the combined active and passive internal bending moment as the ‘bending moment’.

#### 3.1 Optimising internal forces and moments

To calculate the normal force, shear force, and bending moment, we used an optimisation algorithm that determines the best-fitting distributions in space and time. At every point in time, we described the internal forces and moments with a quintic spline^8^ along the length of the fish. Values were prescribed at 12 uniformly spaced control points, between which the values were interpolated with the spline. The first and last control point, at respectively *s* = 0 and *s* = *ℓ*, were fixed at a value of 0, to satisfy the boundary conditions of *N*(0) = *Q*(0) = *M*(0) = 0, and *N*(*ℓ*) = *Q*(*ℓ*) = *M*(*ℓ*) = 0, that should hold for the free ends of a beam.

At a each time step, we optimised the moment-, shear-, and normal distributions to minimise the deviation from equations 7, 8, 9 at every point along the fish. This deviation was quantified by the residual value that is needed to balance the left- and right-hande side of the equations. We optimised the distributions with the Levenberg-Marquardt algorithm^8^, that minimises the squares of the residuals. At every time step, this resulted in a series of control point values describing the internal forces and moments that best satisfy the equations.

#### 3.2 Calculating resultant power

We calculated the resultant power on the fish from two source: the power exerted on the fluid, and the changes in kinetic energy. Both quantities are computed in the inertial reference frame. We calculated the power per unit length that the fish exerts on the water as:

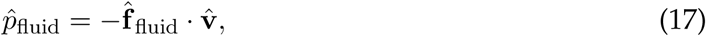

where 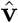 is the velocity of the centreline in world coordinates. We negated the power since we are considering the power that the fish exerts on the water, rather than the inverse.

The kinetic energy per unit length at any point in time was calculated as:

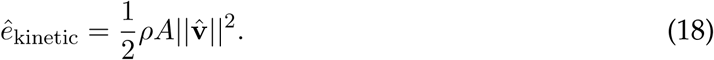

The kinetic power per unit length is the time derivative of the kinetic energy:

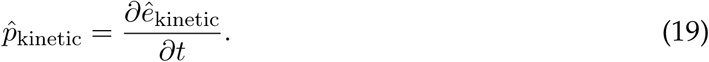

Note that we do not calculate powers based on the internal forces and moments. We do not separate passive and active effects, which might compensate each other. Hence, the power calculated from the net internal forces and moments does not correspond to the actual power consumption by the active components only. This problem is similar to modelling the power consumption of an aeroplane flying at constant speed without separating thrust and drag source: since there are no net forces, the net power calculated is zero. However, when the ‘active’ component (i.e. thrust from the engines) is separated from the ‘passive’ component (i.e. drag of the aircraft), the computed power consumption is clearly non-zero.

### 4 Integrating the equations of motion to generate reference data

We integrated the equations of motion to determine whether the derived equations are physically valid, and to generate reference data to test our algorithm for reconstructing bending moments.

#### 4.1 Constitutive and kinematic relations

The normal forces and moments can be calculated from the displacements using constitutive equations. To generate the reference data, we assumed a Hookean material, resulting in the following equations:

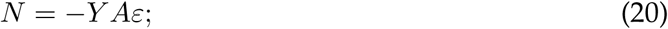

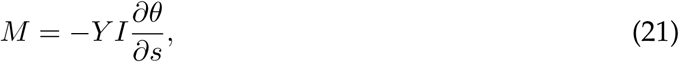

where *Y* is the Young’s modulus. The strain *ε* can be computed from the displacements as follows:

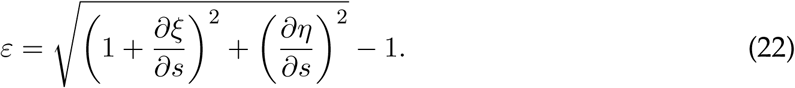

These relations complete the set of equations required to calculate the accelerations of each point on the beam with equations 7 and 8.

#### 4.2 Temporal integration

We used the backward Euler method to integrate the beam accelerations to velocities, and velocities to displacements. To calculate the velocities and displacements in *x*-direction at the time step *i*, we used:

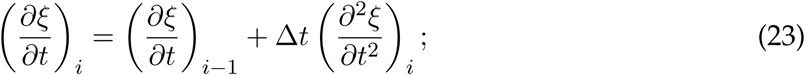

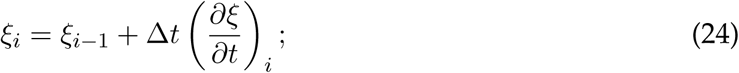

and analogous expressions for *η*.

#### 4.3 Equation scaling

The governing equations resulted in an ill-conditioned system, caused by large scale differences in the matrix coefficients. This makes a system difficult to solve numerically. To improve the condition number, we scaled variables such that all coefficients were close to 1. We used the following scaling coefficients, where an asterisk denotes a scaled quantity:

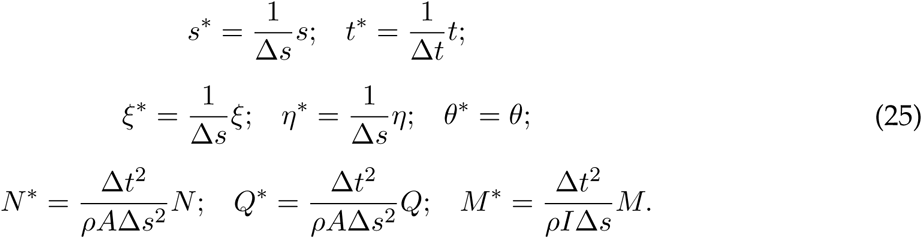

The scaled equations for temporal integration of *ξ* then become:

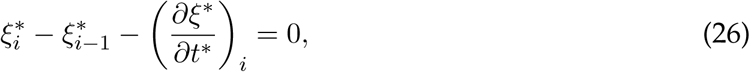

and analogously for the other three equations related to integration of *ξ* and *η*.

Scaling the force balance in *x*-direction yields:

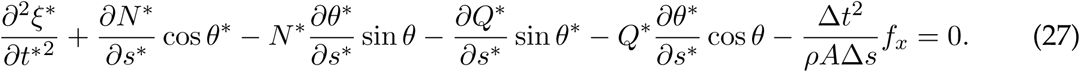

Equivalently, for the force balance in *y*-direction:

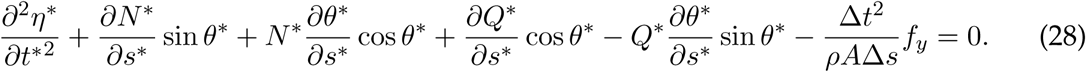

The scaled moment balance becomes:

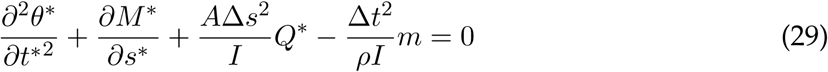

Finally, the constitutive and kinematic relations become:

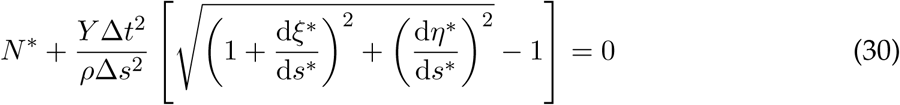

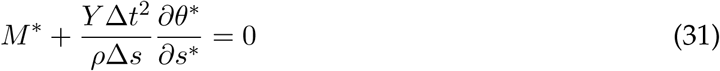

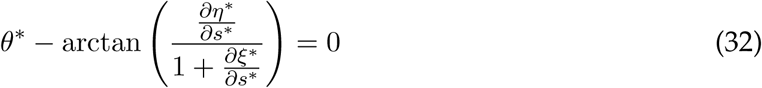

#### 4.4 Solution method

We integrated the scaled equations of motion with multivariate Newton-Raphson, an iterative solution method for non-linear partial differential equations. Based on the solution at the previous time step, we calculated the vector of residuals for each of the 4 temporal integration relations, 3 force and moment balances, and 3 constitutive and kinematics relations per point along the beam. This led to a vector of 10*n*_lon_ residuals, with *n*_lon_ the number of longitudinal points in the beam. To calculate the value for the next iteration, we solved:

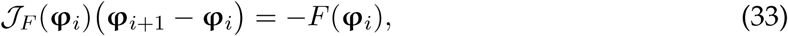

where the subscripts *i* and *i* + 1 denote the current and next iteration, *𝒥*_*F*_ is the Jacobian of the residuals, *F* is the vector of residuals, and **φ** is the vector of variables.

We calculated the Jacobian numerically by perturbing each variable with a fixed-step size and then calculating one-sided finite-differences. We solved the system with a direct solver^9^.

### 5 Validating the inverse dynamics methods

This section describes the validation of the internal forces and moments reconstruction, based on reference data, and based on a reference CFD simulation.

#### 5.1 Internal forces and moment reconstruction

We assessed the validity of the equations of motion by comparing a simulated beam to experimental results^10^. We then tested the reconstruction of internal forces and moments based on reference data produced by integrating the equations of motion (see section 4).

##### 5.1.1 Equations of motion

To assess the validity of the derived beam equations, we compared experimental results from Beléndez et al.^10^ with our simulation of a cantilever beam. In their study, a 300 mm steel ruler (*Y* = 200 GPa) with rectangular cross-section (width × height = 30.4 mm × 0.78 mm) was clamped at one end. The beam was loaded with a point force of 3.92 N in negative *y*-direction at the unclamped tip, in addition to the distributed gravity load of 1.85 N m^−1^ (total 0.554 N over 300 mm).

We simulated a beam with the same geometry and loading, but in a time-dependent manner. We started with the beam in undeformed configuration, then smoothly ramped the loading from 0 to the reference amplitudes over a period of 5 seconds, and continued simulating for 5 more seconds. The resulting deformation was compared to the experimental reference in Fig. 2A—it overlaps strikingly, providing confidence in the physical validity of the derived equations of motion.

##### 5.1.2 Calculating internal forces and moments

To check the correctness of the algorithm for reconstructing forces and moments, we generated a simple model of a ‘swimmer’ by prescribing analytical external forces and moments to a simulated beam of a Hookean material (see section 4). We prescribed the fluid forces as:

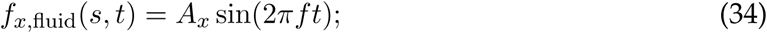

**Fig. 2.**
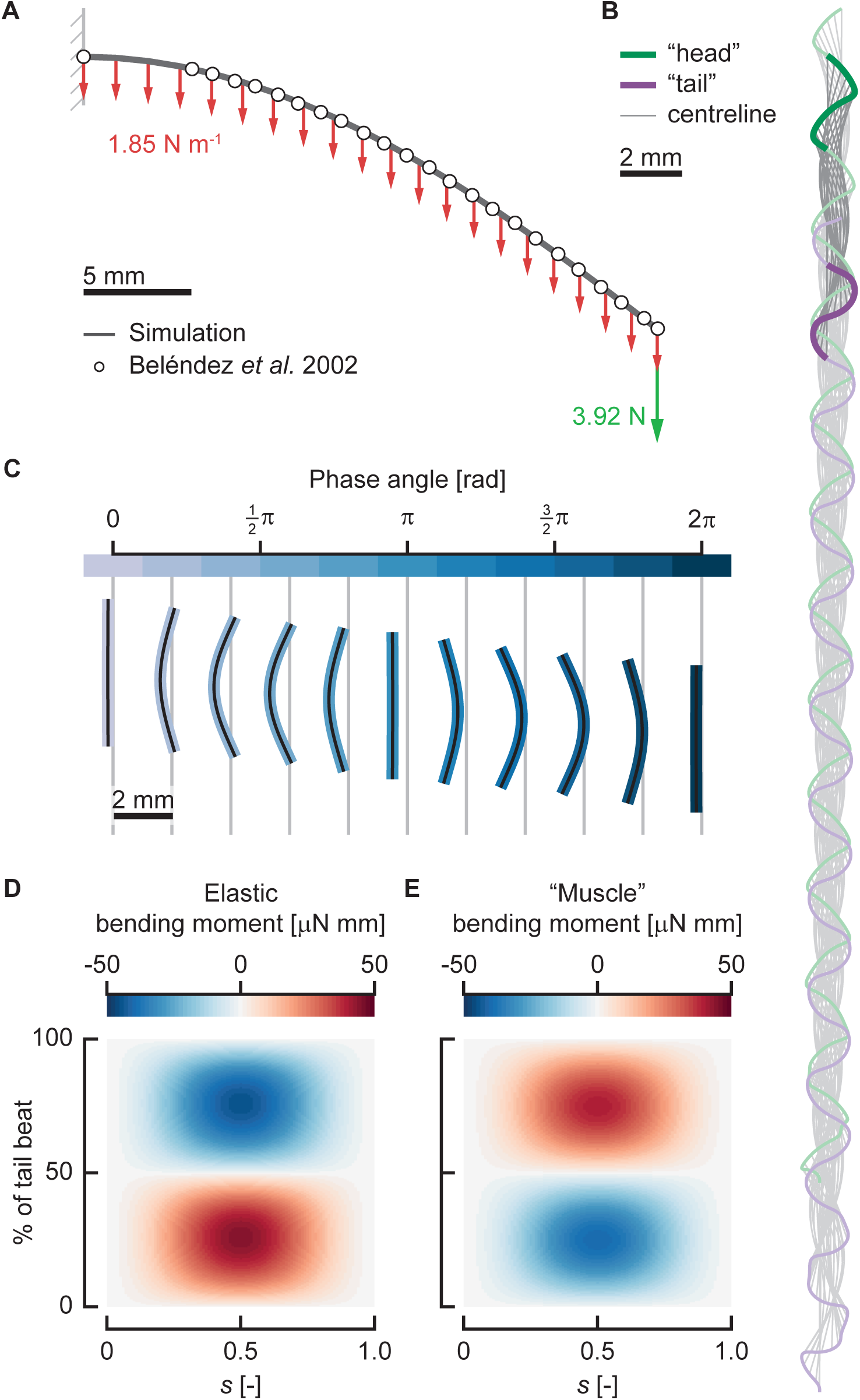
Reference data for the internal forces and moments reconstruction. (A) Comparison of our simulated beam (solid grey line) with experimental results^10^ dots, under a distributed and point load (resp. red and green arrows). (B) Motion of the simulated reference ‘swimmer’, with the beam centrelines (grey), path the ‘head’ (green) and of the ‘tail’ (purple). The dark grey, green, and purple indicate the cycle selected for further analysis. (C) The motion of the reference ‘swimmer’ over the selected cycle, centreline (black), beam coloured by phase angle. (D)–(E) The elastic contribution ((D)) and the ‘muscle’ contribution ((E)) to the internal bending moment for the reference ‘swimmer’.

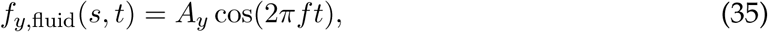

and the muscle moment by:

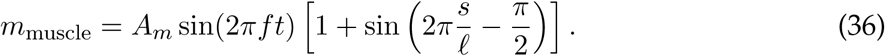

The resulting motion is shown in Fig. 2B, demonstrating a swimming-like motion. We selected a single cycle from the motion (Fig. 2C), after the motion had become reliably periodic after 11 cycles. The total internal bending moment, which is what we reconstruct, consisted of an elastic contribution (Fig. 2D), and a contribution from the ‘muscle’ moment (Fig. 2E).

We reconstructed the total internal forces and moments of the reference ‘swimmer’ with the method described in section 3, see Fig. 3. The results match well, both qualitatively and quantitatively. Qualitatively, the patterns are similar, showing the same dynamics between the reference and the reconstruction. The shear force and normal force show relatively the largest errors (respectively maximum 9.7% and 7.0% of the peak value), while the error in bending moment is low (maximum 0.9% of the peak value). This shows that our main quantity of interest, the bending moment, can be reliably reconstructed with the proposed method.

### 6 Computational fluid dynamics

As a reference to compare the computational fluid dynamics (CFD) results of IBAMR, we used an extensively validated numerical method for fish free swimming^3,4,5^. The curvature of the fish was prescribed similar to Li et al.^4^, by a travelling curvature wave (rather than an amplitude wave in the original reference) with an experimental curvature amplitude envelope of a 3 days post fertilisation zebrafish larva. The frequency was 50 Hz, the fish length 3.8 mm, the water density 1000 kg m^−3^, and the dynamic viscosity 0.8301 mPa s. The motion of the fish was calculated based on the fluid dynamic forces, resulting in a free-swimming fish. We used the resulting fluid force distributions and motion to calculate internal forces and moments.

We prescribed the same motion with a custom-developed add-on to the open-source immersed boundary method IBAMR^6^, see section 2. Note that we did not integrate the motion of the fish, but prescribed the position of the fish surface directly at all time points. For the results from IBAMR, we also calculated the internal forces and moments to compare to the reference from the validated solver.

**Fig. 3.**
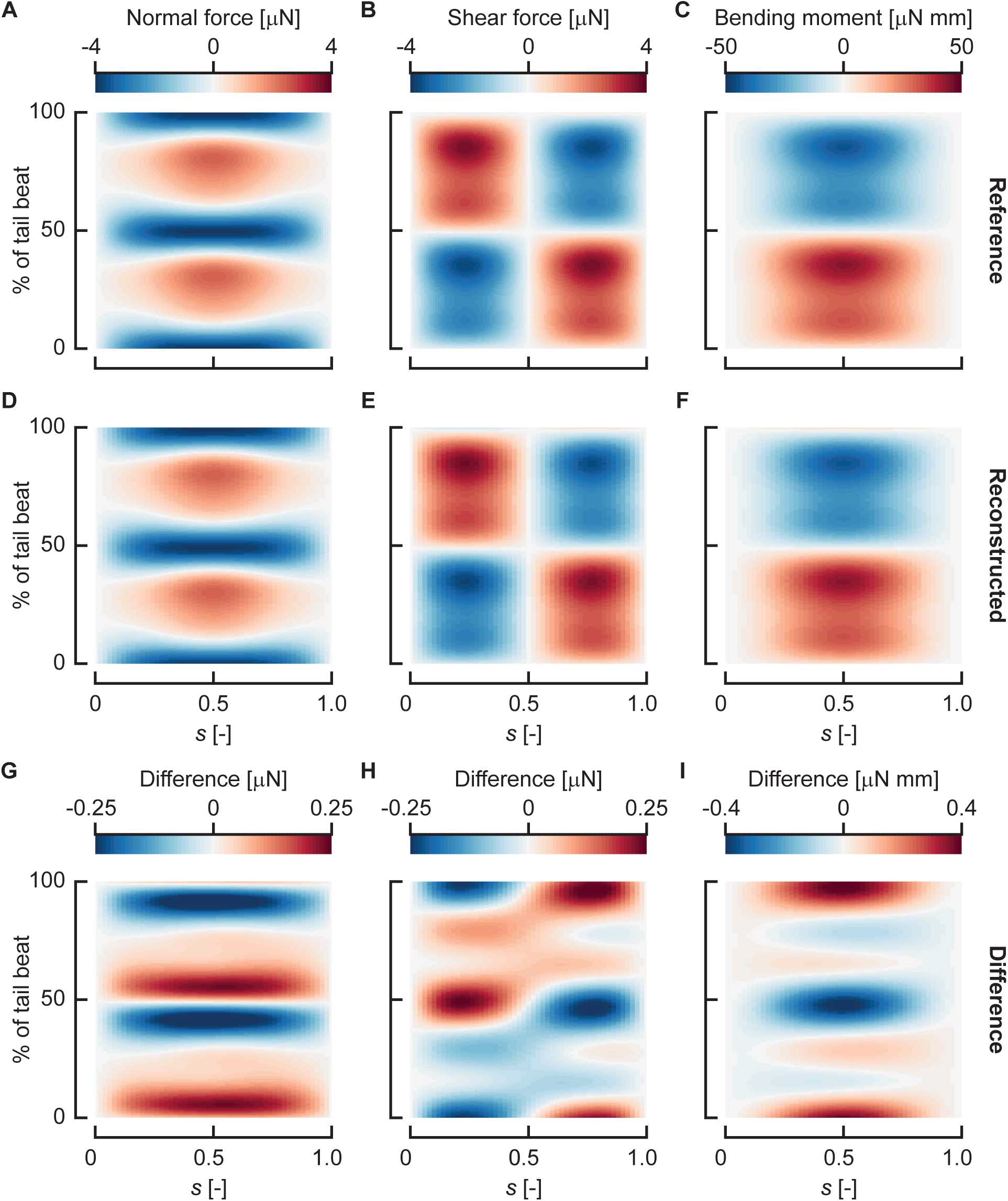
Comparison between reference and reconstructed internal forces and moments. (A)–(C) Reference normal force ((A)), shear force ((B)), and bending moment ((C)). (D)–(F) Reconstructed normal force ((D)), shear force ((E)), and bending moment ((F)). (G)–(I) Difference between the reference and the reconstructed normal force ((G)), shear force ((H)), and bending moment ((I)).

#### 6.1 Influence of the surface offset

We calculated the force distributions on the fish by interpolating quantities from the CFD flow field to a triangulated surface of slightly offset from the skin of the fish. The amplitude of these forces is dependent on this offset, related to the accuracy of the interpolation of the pressure and velocity gradients. Due to the immersed boundary approach, there is also a flow field inside the fish, but this is not physically relevant for the force calculations. If this is taken into account in the interpolation, errors in the force distribution will occur.

Fig. 4A shows the effect of the surface offset on the accuracy of the bending moment reconstruction. The optimal distance for the offset surface for a simulation with a finest mesh size of 15 µm (the final selected mesh size) was found to be 20 µm. This distance guaranteed that the flow field was interpolated from only cells outside the body, but was close enough to accurately reconstruct the frictional forces.

**Fig. 4.**
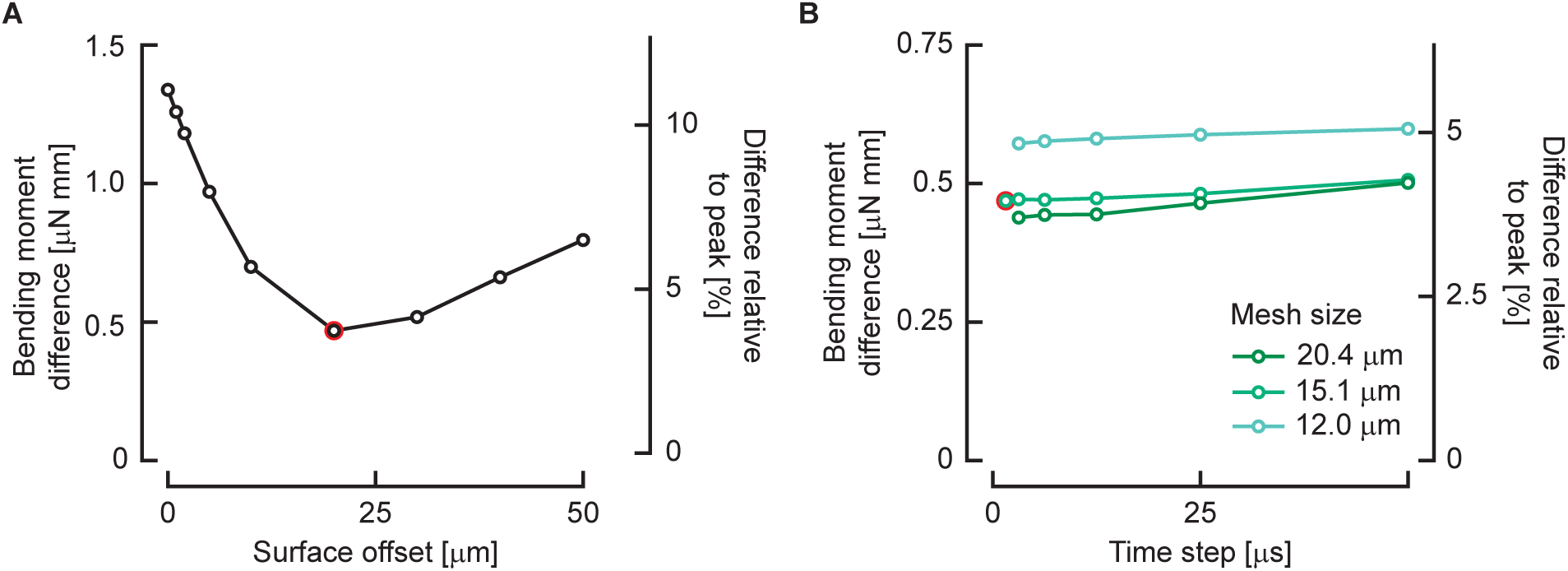
Convergence of the immersed boundary solver. (A)–(B) The difference in reconstructed bending moment compared to the reference solution as a function of the offset of the surface used for force calculations ((A)), and the mesh size and time step ((B)). The final choices of surface offset, mesh size, and time step size are highlighted with a red circle.

#### 6.2 Influence of the mesh size and time step

To assess the dependency of the solution on the mesh size and time step, we simulated the reference case from the validated solver on three different mesh sizes (finest level size 20.4, 15.1, 12.0 µm), and 5–6 different time steps. The time steps were chosen such that the coarsest step always led to a maximum CFL number close to (but below) 1. In the tested range, mesh size nor time step had a large influence on the solution.

For mesh size, the largest step in accuracy is from 20.4 µm to 15.1 µm, the step to 12.0 µm is smaller. We chose a mesh size of 15.1 µm, as it allows reasonable accuracy while remaining computationally feasible—memory usage is a limiting factor on our computational facilities as the meshes get larger. For the time step, smaller time steps lead to marginally smaller errors. Computation times are at most linearly affected by the time step, so the trade-off for computational feasiblity is less relevant. Hence, to be on the safe side, we chose a time step of µs for solving the fluid dynamics.

A comparison of the reference simulation with the final choice of mesh size 15.1 µm and time step 0.5 µs is shown in Fig. 5. The bending moments show qualitatively similar patterns, in time and space. The magnitude is slightly underestimated in IBAMR compared to the reference solution.

**Fig. 5.**
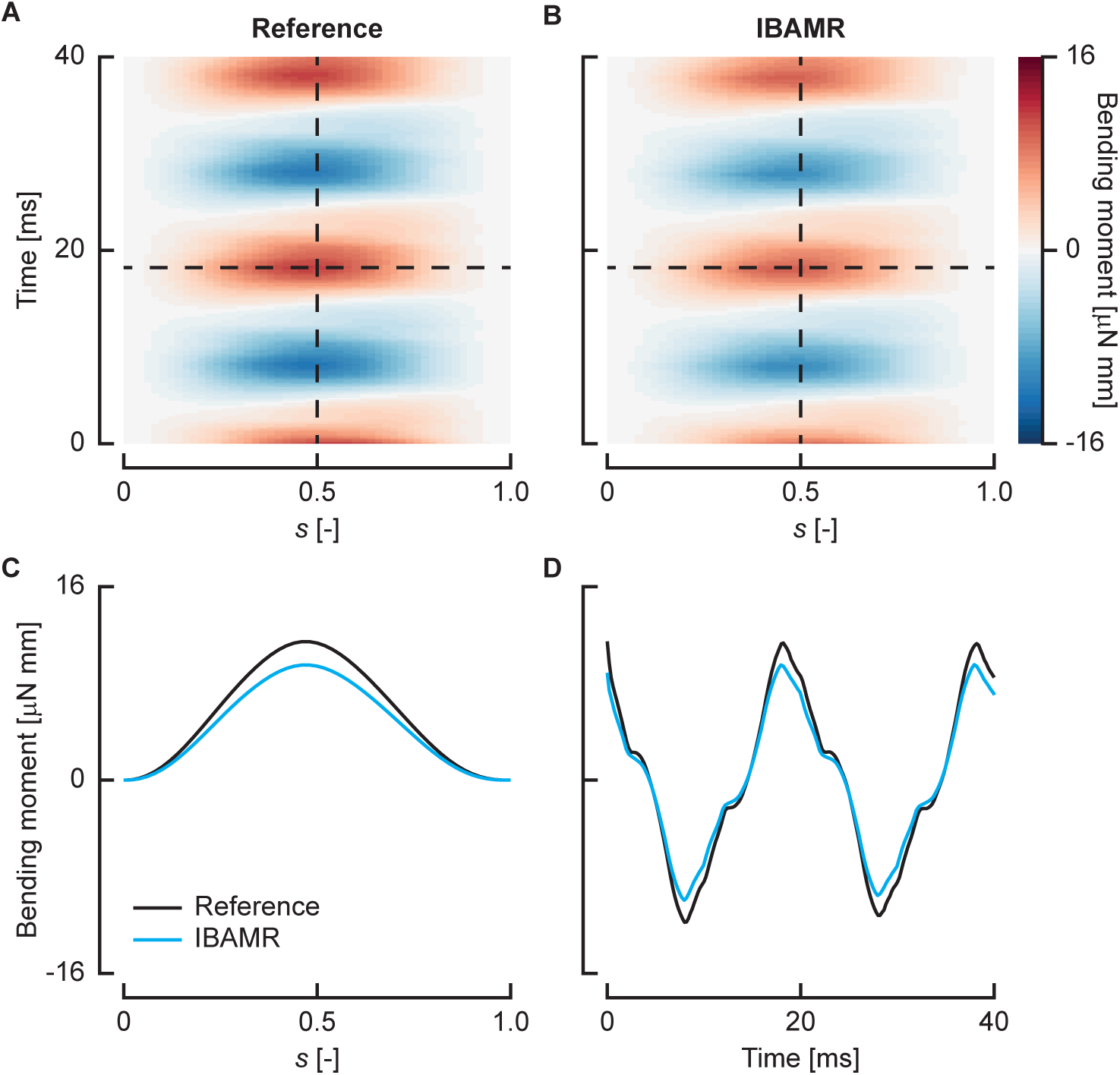
Comparison between the validated solver and the final choice of immersed boundary solver settings. (A)–(B) The reconstructed bending moment for the fluid dynamic forces from the reference solver (A) and IBAMR (B). The dashed lines indicate the two slices shown in (C) and (D), at a single time step, and a single position along the fish.

